# Bone Surface Mapping: a new method for recording detailed human bone surface variation

**DOI:** 10.1101/2022.06.21.496497

**Authors:** Sarah E. Paris, Robert A. Foley

## Abstract

This paper presents a new method for mapping surface features on bone (e.g. presence and distribution of ochre, surface cuts and scratches, pathologies, etc.). It is an accessible and quantifiable alternative to photogrammetry or three-dimensional scanning. It breaks each bone down into a number of two-dimensional geometric faces that can be utilised in statistical analyses. The number of faces differs depending on the skeletal development of the individual, with children having fewer faces. The recording method allows faces to be grouped by anatomical plane, bone type, side, and region of the body. These selective groupings facilitate flexible categories for hypothesis testing. The technique was developed to enable a detailed recording of ochre staining on human skeletal remains from Khok Phanom Di, Thailand, used in conjunction with a novel recording method to quantify pigment use across the body, and some exemplary results are presented here. While the technique was developed for recording bone surface staining it could also be used to map other variations on bone surfaces, such as in palaeopathological, forensic or taphonomic contexts.

## Introduction

There are many methods and standards for recording the human skeleton outlined in multiple osteo-disciplines, for example: field archaeology, osteology, bioarchaeology, palaeopathology, biological anthropology, and forensic anthropology (Brothwell, 1994; Buikstra and Ubelaker, 1994; Austin and King, 2016; Brickley, 2017). Each discipline has different purposes and requirements and so the existing recording methods vary from a visual representation of a skeleton to a tick-list of skeletal elements, or a combination. Buikstra and Ubelaker (1994) remains one of the most thorough collections of documenting methods, with 56 pages for recording human remains. Their work covers isolated individuals to comingled remains; bones and teeth of adults, children, and infants; osteological information relating to sex, age, trauma, and pathology; burials and cremations; and taphonomy. The section on taphonomy includes “other cultural modifications”, leaving space for bone type, location, photographs/drawings, and comments (Buikstra and Ubelaker, 1994, Attachment 24). As part of skeletal recording, osteo-disciplines regularly have to record patterns on the surface of bones. In forensics or palaeopathology skeletal mapping is often a diagnostic tool to identify areas where trauma or unusual bone formation occurs. Figure 1, adapted from Lewis (2000), shows the use of skeletal mapping to illustrate the ‘distribution of lesions on the infant skeleton as a result of metabolic or infectious disease’ (51). This visual representation of osteological changes is effective; however, it normally denotes the ‘abnormal’ within an assemblage, thus there is little need for the method to be particularly comparative, unless looking at the root cause across multiple groups. Through necessity, forensics is at the forefront of these types of comparative analyses as there is a need to provide evidence for repeated, sustained, or characteristic trauma. Novak’s (2006) work on skeletal trauma and domestic violence provides a framework by which individuals can be compared through statistical analysis by breaking down the human skeleton into 54 anatomical areas and zones. However, for some purposes this is not sufficiently detailed. In this vein, the method reported here broke the skeleton down into anatomical zones (skulls, upper limb, hands, torso, pelvis, lower limbs, and feet), further broken down into individual bones, side of the body (including central), and their anatomical planes. This allowed for a more in-depth exploration of surface trait distribution, originally for ochre, but adaptable for other purposes.

**Figure 1.**
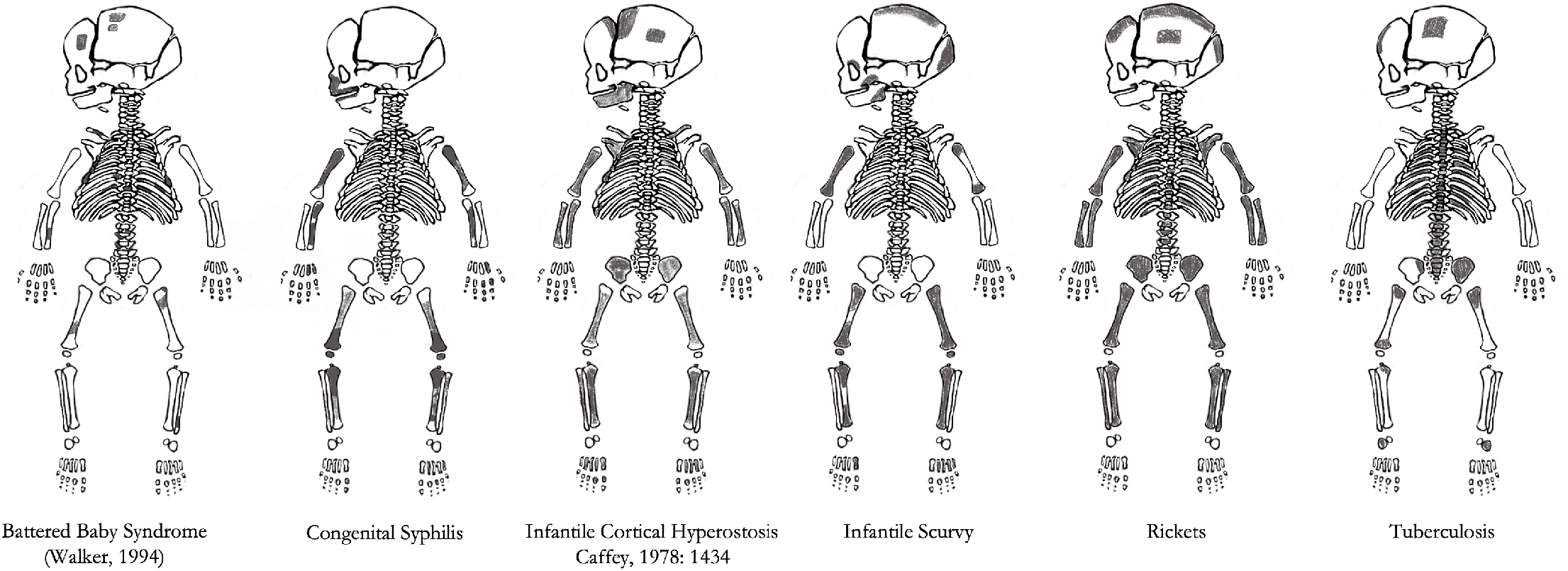
“Distribution of lesions on the non-adult skeleton as a result of metabolic or infectious disease” (adapted from Lewis, 2000, 51). The shading indicates areas of abnormal bone formation.

## The method

### Adults

By visualising each bone as a simple three-dimensional geometric shape that can be folded out into a net (Figure 2a), the bone surfaces of the skeleton were divided into 521 two-dimensional faces for adults, with an adjusted number for sub-adults based on skeletal development (Figure 2b; supplementary materials). Determining the faces of each net was by considering each bone as a three-dimensional geometric shape with the fewest number of sides in order to capture the dominant features of the bone. The starting point for each long bone was to consider it as a cuboid with six faces made up on the anterior, posterior, inferior, superior, medial and lateral anatomical planes; geometric faces were added to capture the morphological characteristics of each bone. For example, the 24 femoral faces comprise the visible anatomical planes of the neck, trochanter, head, shaft and distal features (as illustrated in Figure X). For post cranial axial skeleton, the faces were simplified due to the blade like morphology of the ribs, corpus sterni and manubrium. These bones only have anterior and posterior (sternum) or internal and external faces (ribs). The ribs were further simplified by only distinguishing the first rib, grouping the second to twelfth ribs. This decision was made to simplify identification and mitigate preservation issues; however, by adding an additional twenty faces (left and right), to the existing eight the method can be adapted to suit the needs of any research question. Each vertebra was considered in the six anatomical planes. The carpals, tarsals and metapodials were simplified to faces in the dorsal and palmer/planter plane for the left and right. The phalanges were distinguished by hand and foot; proximal, intermediate, and distal; the dorsal and palmer/planter plane; but not the left and right. The cranium consists of 54 planes, recording the visible face of each of the constituent bones in each anatomical plane (palate, sphenoid, temporal, occipital, parietal, frontal, zygomatic, maxilla), and two faces for the collective bones of the left and right orbits. The cranium includes internal and external planes as the method was devised for archaeological collections where skulls are frequently broken; however, these should be recorded as absent where a skull is complete or omitted if all skulls in the analysis are complete. The faces of the mandible separate the condyle, coronoid, ramus, gonial, border, and body across the visible anatomical planes. Teeth are listed collectively as either maxillary or mandibular, with no distinction made between buccal/labial or lingual (again, these could be added as the user requires).

**Figure 2.**
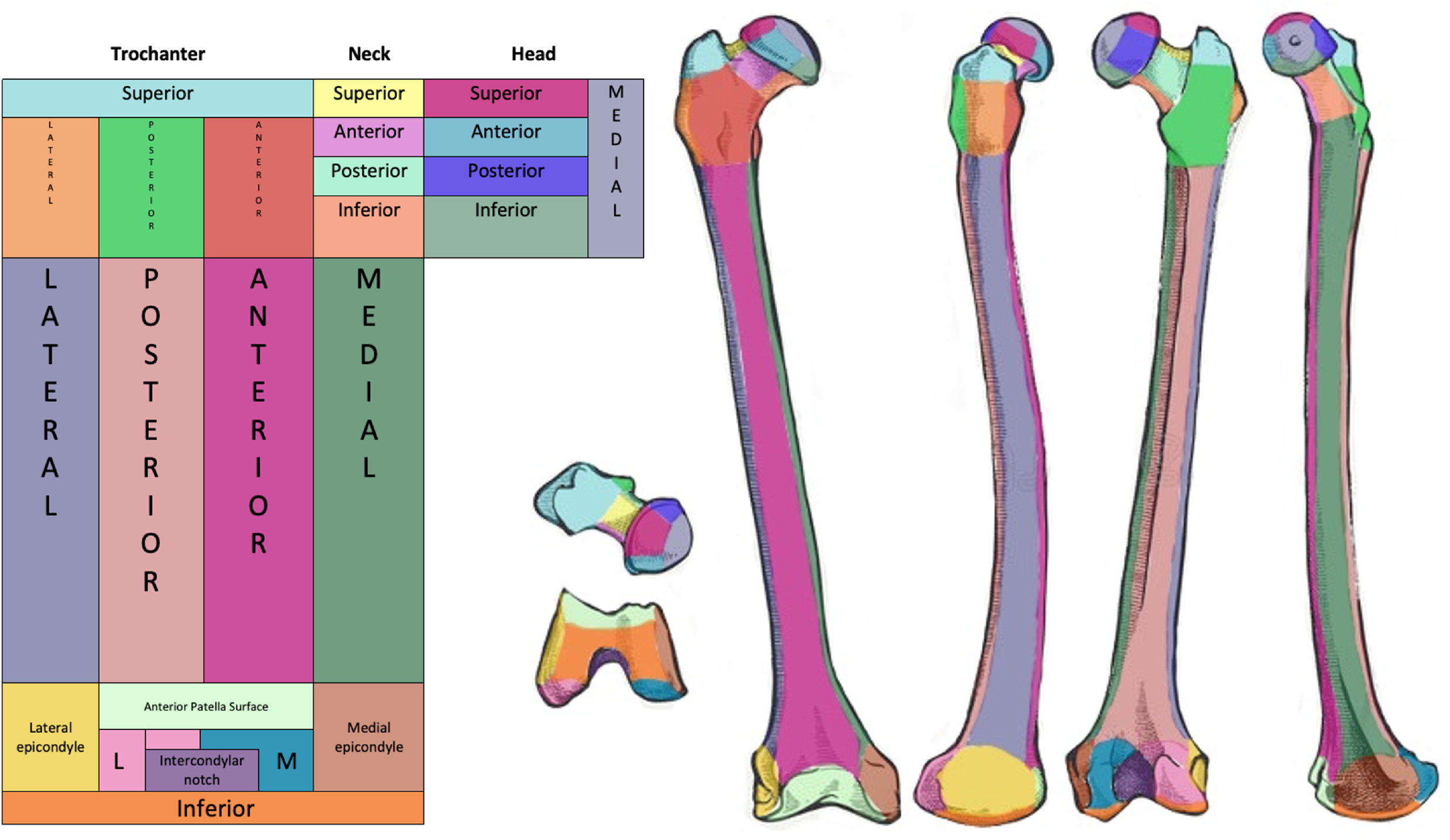
Example of bone mapping method, right femur. a.) Two-dimensional net of the femoral bone faces (left). b.) Three-dimensional application of bone mapping regions onto the right femur. Corresponding regions are identified using matching colours.

### Sub-adults

Juveniles could arguably have more faces as there is a separation of unfused epiphyses and their corresponding growth plates, this creates more visible surfaces. These surfaces could have been added to the method; however, in practice this would not be practical as developmental changes have a window, some more sex determined than others, thus it is impossible to predict exactly which surfaces should be present. Additional complications are preservation, collection, and identification bias. Ossification centres may have been present during life but due to their small size could have been missed during excavation, not preserved, miss-identified, or scavenged. In order to maintain the skeletal regional categories for comparative analysis of juveniles with adults the corresponding area of the bone at the site of epiphyseal fusion was used. The epiphyses themselves were only considered when they were sufficiently formed to be identifiable. Scheuer and Black (2004) describe juvenile development by each bone in extensive detail; they not only list the estimated age ranges for completion, but they also differentiate between visible ossification centres and point of distinct development specifically for the purpose of identification. The maximum developmental age was taken from Scheuer and Black (2004) to enable a record of completeness. By eliminating planes that would be unidentifiable at the estimated age for the individual and using the correlating adult regions, the number of bone faces for juveniles was lower than adults, varying at different stages of development (Figure 3).

**Figure 3.**
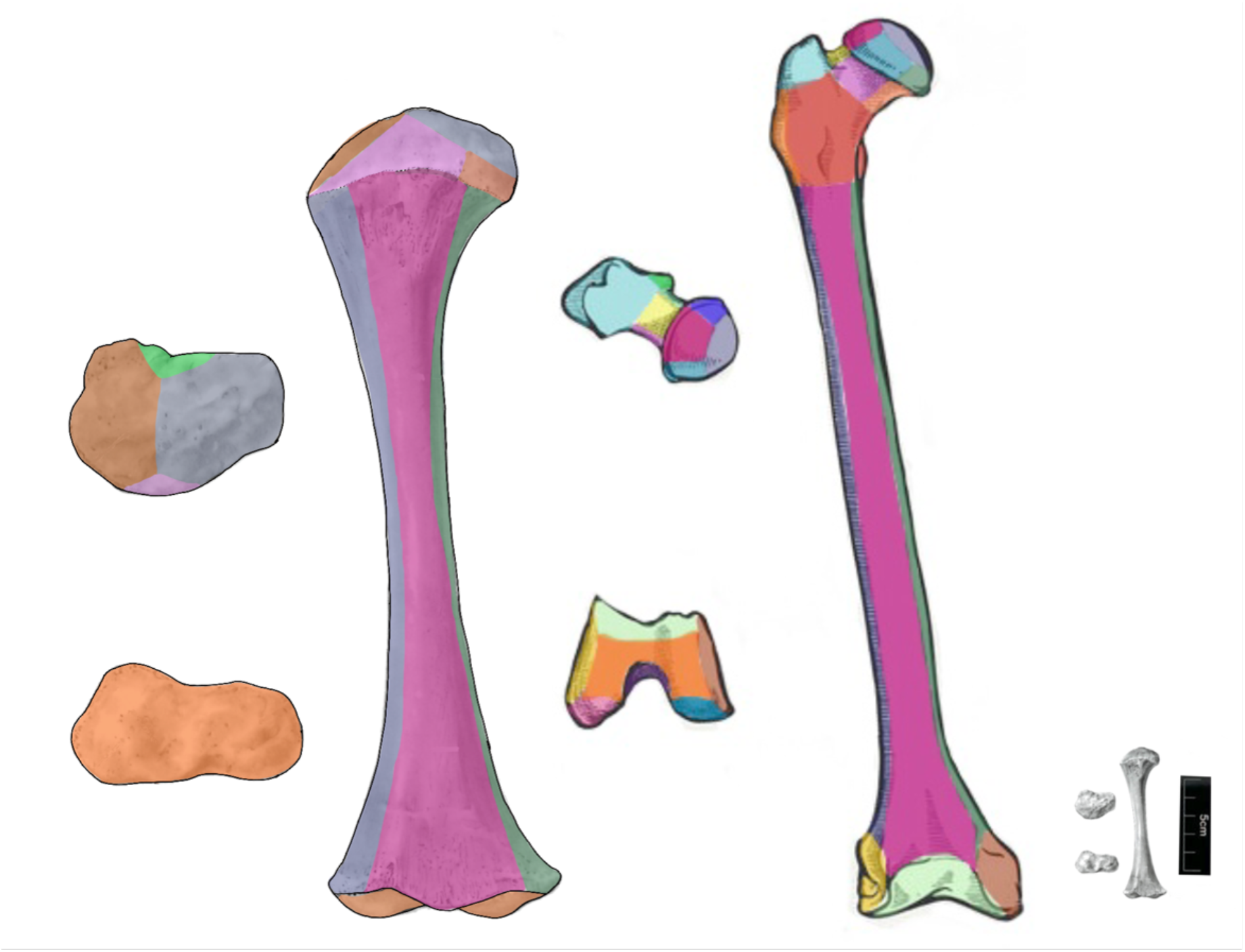
Comparative faces on a perinate (enlarged, left) and adult (right) right femora. The greyscale juvenile femur (bottom right) is included for approximate scale with the adult femur (scale measures 5cm).

### Nomenclature

The faces were given unique five character names or ‘codes’ based on anatomical features, there were: ‘region’, ‘bone’, ‘specific’ (or descriptor), ‘side’, and ‘anatomical plane’, which can be used to group faces for analysis. The ‘code’ for each face is the first letter of each category. For example, the face corresponding to the anterior femoral shaft of the right leg would be LFSRA (lower limb, femur, shaft, right, anterior). Where this would create a duplicate code the ‘bone’ or ‘specific’ letter was substituted for a unique letter. For example, the lower limb, femur, trochanter, right, anterior would be the same as the lower limb, femur, trochlea, right, anterior: LFTRA. So, the former keeps the original code, and the trochlea is listed as A_trochlea on the data collection sheet with the code LFARA.

### Boundary issues

Clearly, as bones comprise continuous surfaces, the faces are to some extent arbitrary units along this continuum. To minimise inconsistencies, all bones were oriented anterior-posteriorly, and then rotated through 90 degrees to obtain the four faces and similarly, superior-inferiorly and rotated 180 degrees to obtain the faces in the fifth and sixth anatomical plane. Where the observable trait continues across faces, it is recorded on each face where it occurs (Figure 4). The boundaries provide a flexible framework through which hypotheses can be tested comparing human remains on an individual scale and across populations.

**Figure 4.**
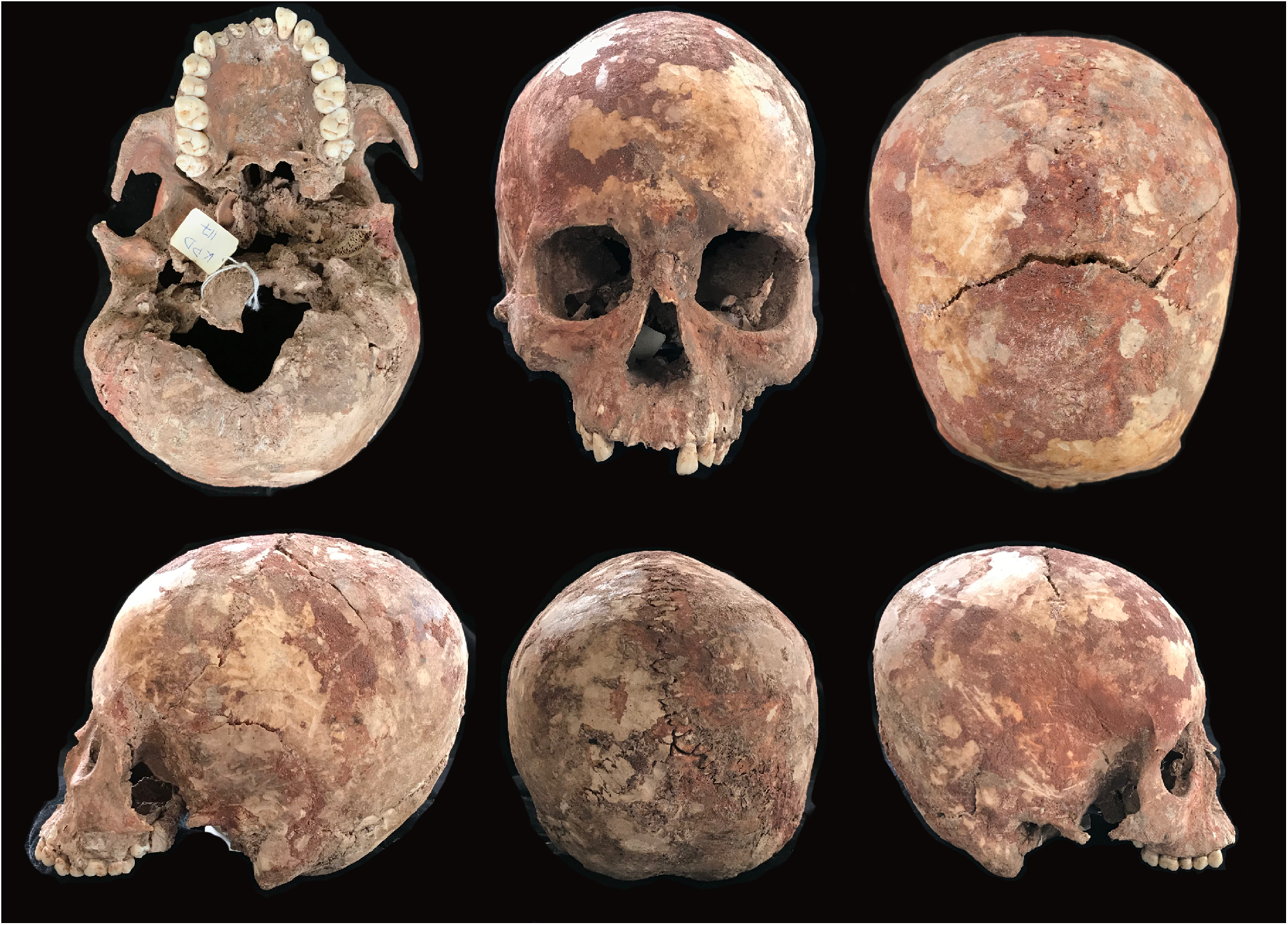
Khok Phanom Di Burial 117 skull. The skull is shown in the inferior, anterior, superior, left lateral, posterior, right lateral orientation (from top left to bottom right). Each rotation enables the recording of the faces associated with that anatomical plane.

### Preservation and incompleteness

The amount of surface staining on a skeleton is inevitably affected by preservation and the amount of remaining bone per individual. Thus, any method recording skeletal staining must account for the completeness and preservation of each individual. What exactly is recorded for each geometric face can be decided based on the granular need of the research question. It could be a quite simple binary presence/absence, a percentage of face covering, or a more complicated scale. Analysing the data will again come down to the specific research question and the size of the data set.

### Analysis

The individual faces can be compared between individuals using the ‘codes’ listed for each face (explained in more detail below); however, anatomical features can be compared based on specific research questions by grouping together specific faces. These can be determined by the user or for ease of analysis they can be grouped using one of the four categories established in the recording sheet: ‘region’, ‘bone’, ‘side’, and ‘anatomical plane’. The ‘region’ comprises of eleven regions, and within those 57 ‘bones’. Each face has a ‘side’, left, right or centre (with the addition of hand and foot for the phalanges). The six ‘anatomical planes’ are included with the addition of palmer/planter and dorsal for the hands and feet; and internal and external for the ribs. An excel spreadsheet with the list of bone planes and their categories is included in the supplementary materials for ease of use. By having the anatomical categories as columns built into the recording sheet it enables comparison between sides of the body as well as individual bones. For example, if there was a significant correlation between two bony elements, for example staining on proximal and distal posterior femurs, this could be considered against the femoral anatomy, which might lead to an interpretation that staining is more prevalent on faces which are in contact with the ground.

In addition to the code and five anatomical categories the developmental age of each the feature of each face is also included. This provides an analytical tool to remove faces based on skeletal age but does require an age assessment of the skeleton based on the same ageing methods used here (Scheuer and Black, 2004). By separating out the five aspects of the faces when undertaking analysis, it is easier to single out these categories for comparison.

## Results

### An example: Ochre Recording for Individuals from Khok Phanom Di

The methodology described above was applied to record ochre staining on 152 individuals from the Khok Phanom Di, an agricultural transition site from central Thailand (Higham 1989; Higham and Bannanurag, 1990; 1991; Higham and Thosarat, 1993; 2004; Thompson and Higham, 1996; Tayles, 1999; Vincent, 2004). Two data sets were created, the first a simple binary presence or absence of ochre, then a more complex series of ochre scores, which considered the area (scale 0-7), density (scale 0-6), and thickness (scale 0-5) of the pigment (Paris 2021, Paris and Foley in prep). Damage to the bone surface was also recorded where this obscured the pigment, in a fourth category with a zero to three scale. Where bones or faces were absent or obscured, they were recorded with an “A” to differentiate from a score of 0, which would imply no ochre visible on the bone. Where a bone face was partially represented a score was still given, thus the completeness of each skeleton as represented by ochre scores can be higher than the percentage of the skeleton present.

Despite the high standard of preservation and the presence of small bones (for example the hyoid and phalanges), which are often missed during excavation, it was not always possible to identify small fragments of bone (Daniell, 1996; Lewis, 2000; Cunningham, et al., 2016). Any bones that could not be determined or sided were excluded from the study as without identification these bones could not contribute to the data on ochre distribution. In addition, ossification centres in their earliest stages “are indistinguishable from each other, they are identified by their anatomical position rather than their distinctive morphology. They therefore require the presence of soft tissue to hold them in place to allow identity to be established” (Scheuer and Black, 2004, 14). This is a frequent problem in archaeological contexts where no soft tissue remains, and the individual bones are not identified and kept separately at point of recovery. The difficulty of correctly attributing and siding epiphyses and hyoids of younger individuals correctly meant that these elements were routinely excluded from the study for younger individuals.

We can show the power of the method by briefly describing patterns of variation in ochre presence-absence across the skeletal faces. Figure 5 is a visual representation of the presence or absence of pigment for the Khok Phanom Di assemblage. The 124 individuals from the site who had ochre recorded on their skeleton listed in the y-axis and 424 bone faces selected for comparison (based on preservation bias) in the x-axis. The red tiles demonstrate faces which had ochre present and the grey those where no pigment was observed. The white tiles demonstrate where the bone was absent. The faces are ordered broadly anatomically, with the skull on the left and feet on the right. The individuals are ordered by the estimated biological age of the individual, with the youngest at the bottom and the oldest at the top. The preservation bias and developmental bone absences can be seen more frequently at the bottom of the graph, amongst the youngest individuals.

**Figure 5.**
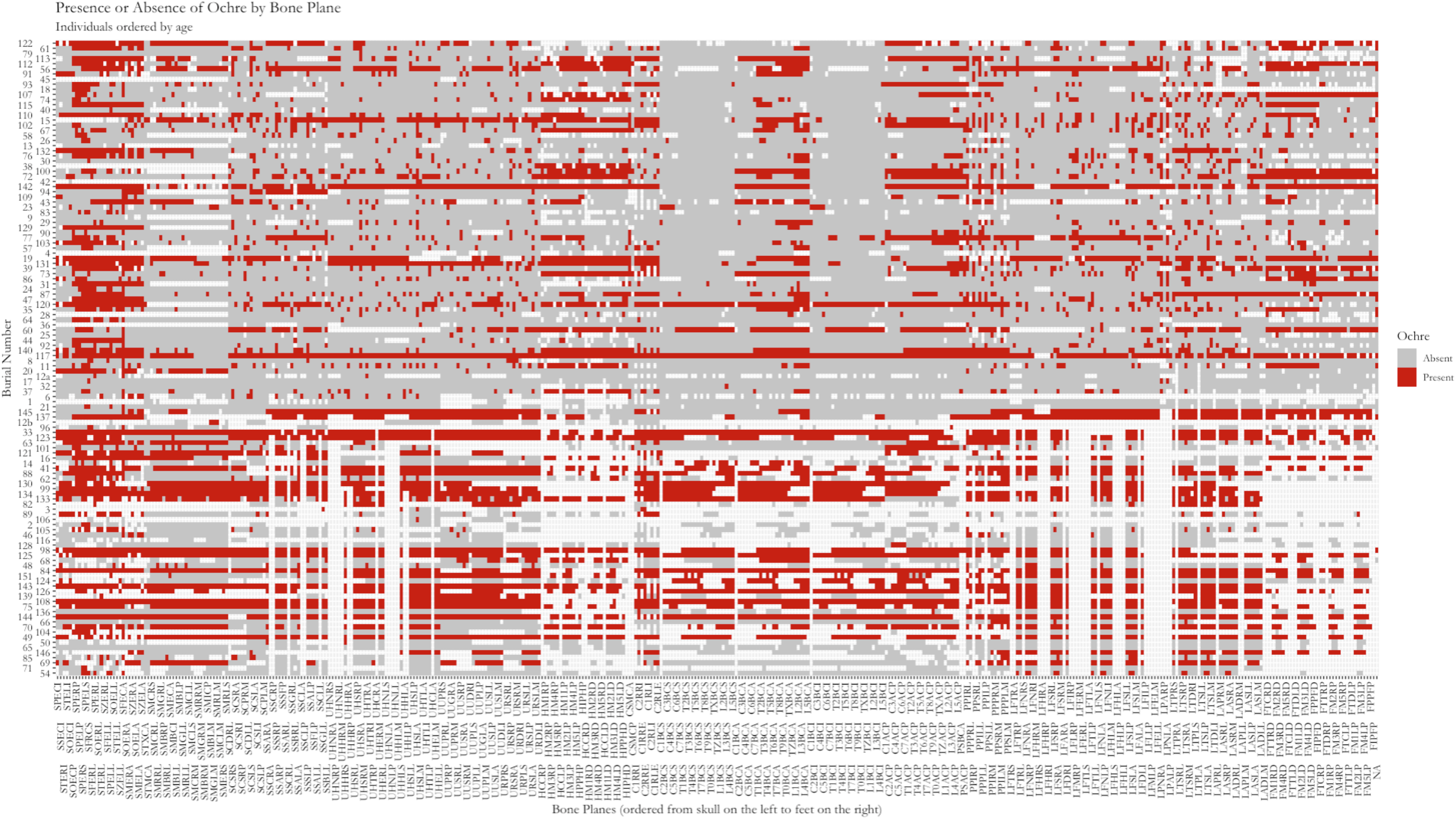
The presence or absence of ochre across 424 bone planes per individual. Individuals ordered by age, youngest at the bottom; bone planes ordered head to toe, skull on the left. Red tiles indicate planes with ochre, grey those without pigment, white tiles show missing planes.

The data can be partitioned easily to explore frequency patterns. For example, we can compare the frequency of ochre anterior and posterior planes of the humerus and femora. Figure 6 shows that the majority of faces on both the femora and humeri do not have pigment present. The proportion of ochred to non-ochred varies marginally between the two bone types and anatomical planes. The two bone types differ in which anatomical planes are more commonly observed to have ochre.

**Figure 6.**
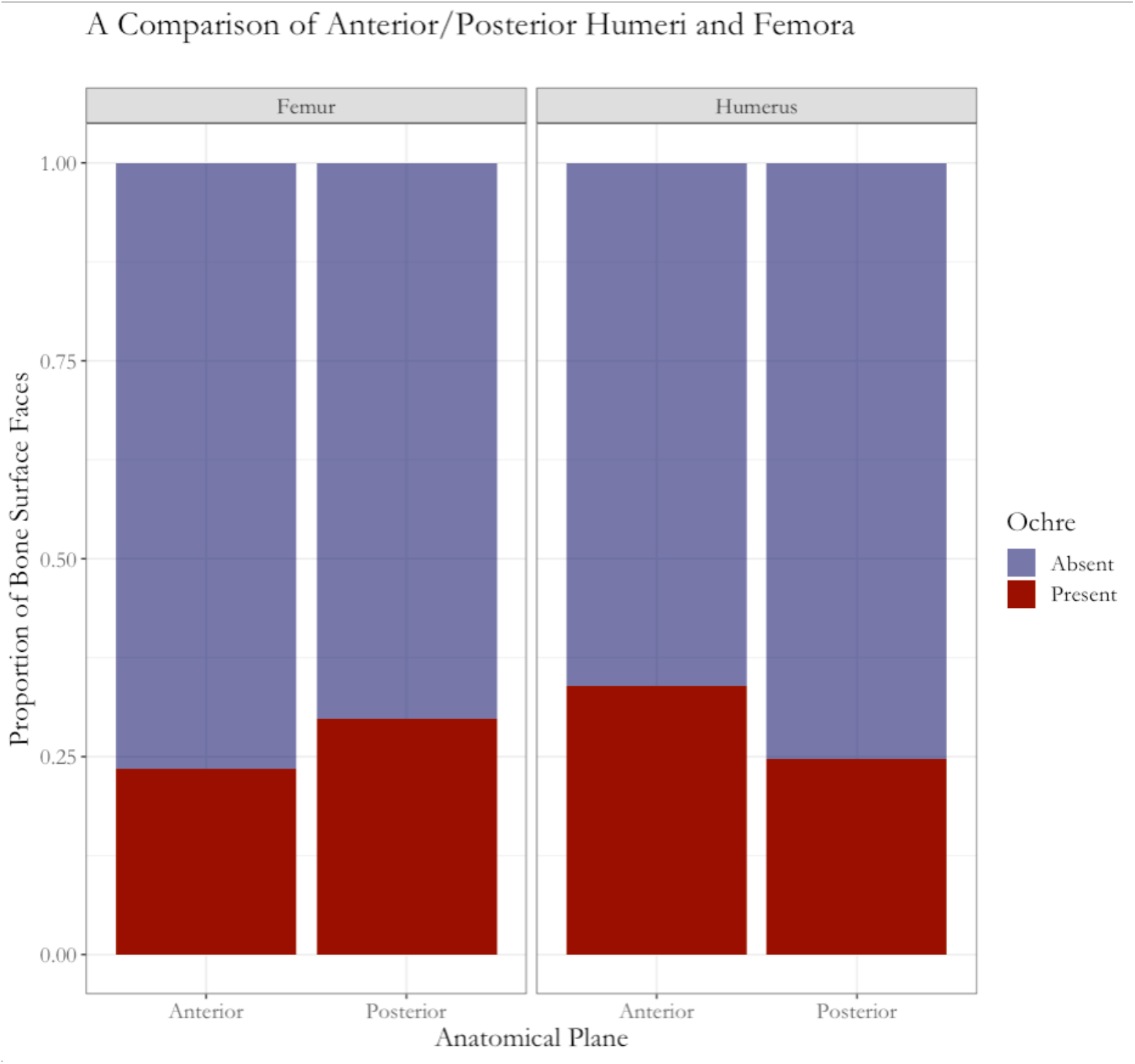
A comparison of the frequency of ochre presence and absence on the anterior and posterior planes of the humerus and femora.

Once the data have been organised, they can also be analysed in terms of their context – for example, do patterns vary by age or sex? Figure 7 illustrates that the percentage of faces per individual with ochre present varies cross both features. All three groups have a large proportion of individuals clustering at the lower end of percentage ochre coverage. Males have fewer individuals with no pigment than females and juveniles. Juveniles differ from adults with a large proportion of individuals having a high percentage of ochre coverage.

**Figure 7.**
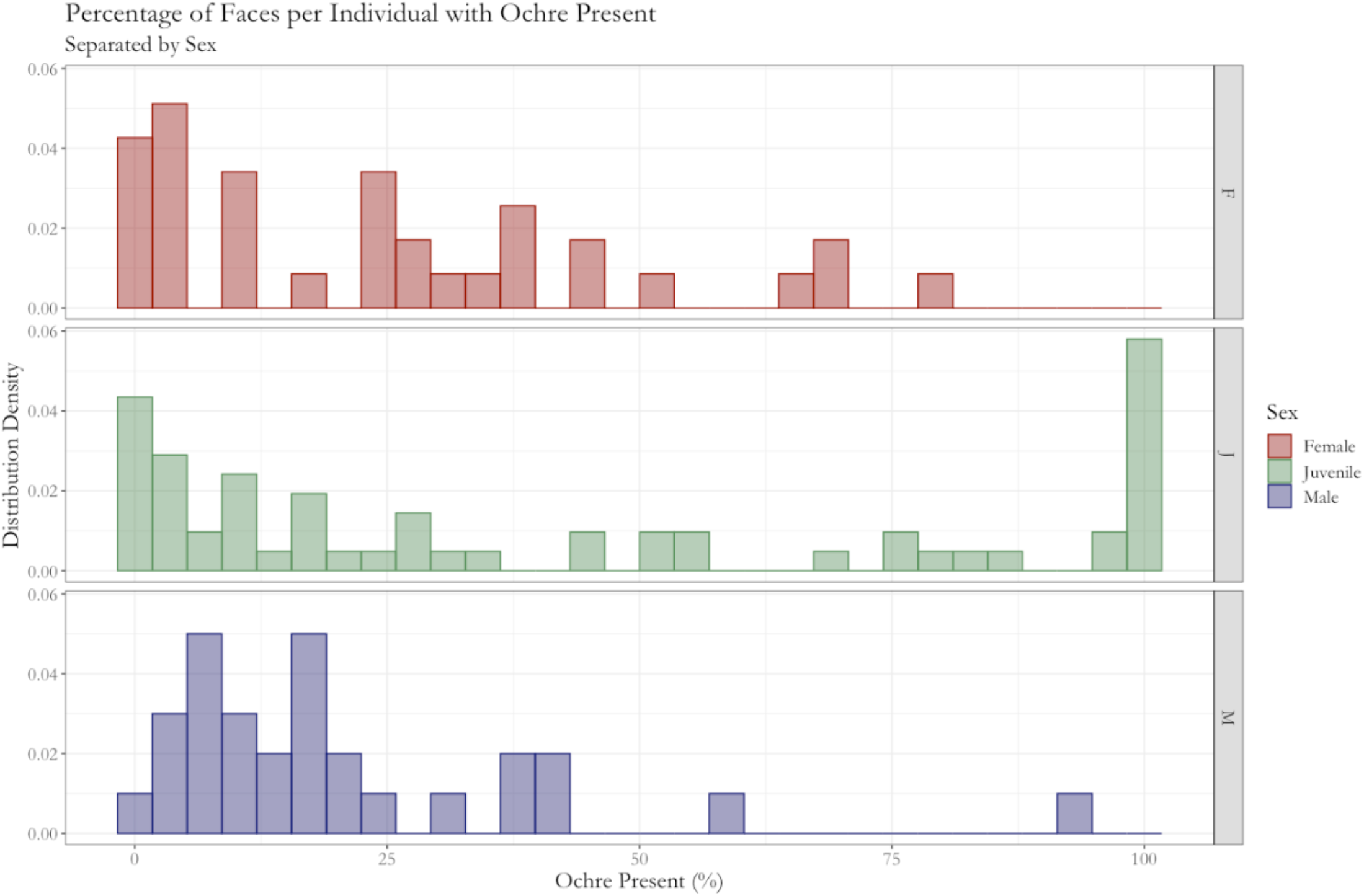
The percentage of faces with ochre present per individual, separating juveniles and sub-adults, with adults further divided by sex. The variation across the three groups demonstrates different patterns of ochre use across osteobiographical factors.

Once initial patterns have me observed anatomically or osteobiographically they can be explored in combination. At Khok Phanom Di the most dominant osteobiographical feature influencing pigment use was age. Adults had a greater level of bodily variation of pigment distribution than children. This was explored further through factor analysis. Through this a heatmap was created demonstrating the percentage of individuals with ochre across the pigmented adult assemblage (Figure 8). The heatmap clearly demonstrates that bodily ochre distribution was more varied, while the skull (excluding the posterior) had a high proportion of pigment coverage across the majority of adults. The significance of these results is discussed in greater detail in Paris (2021; in prep).

**Figure 8.**
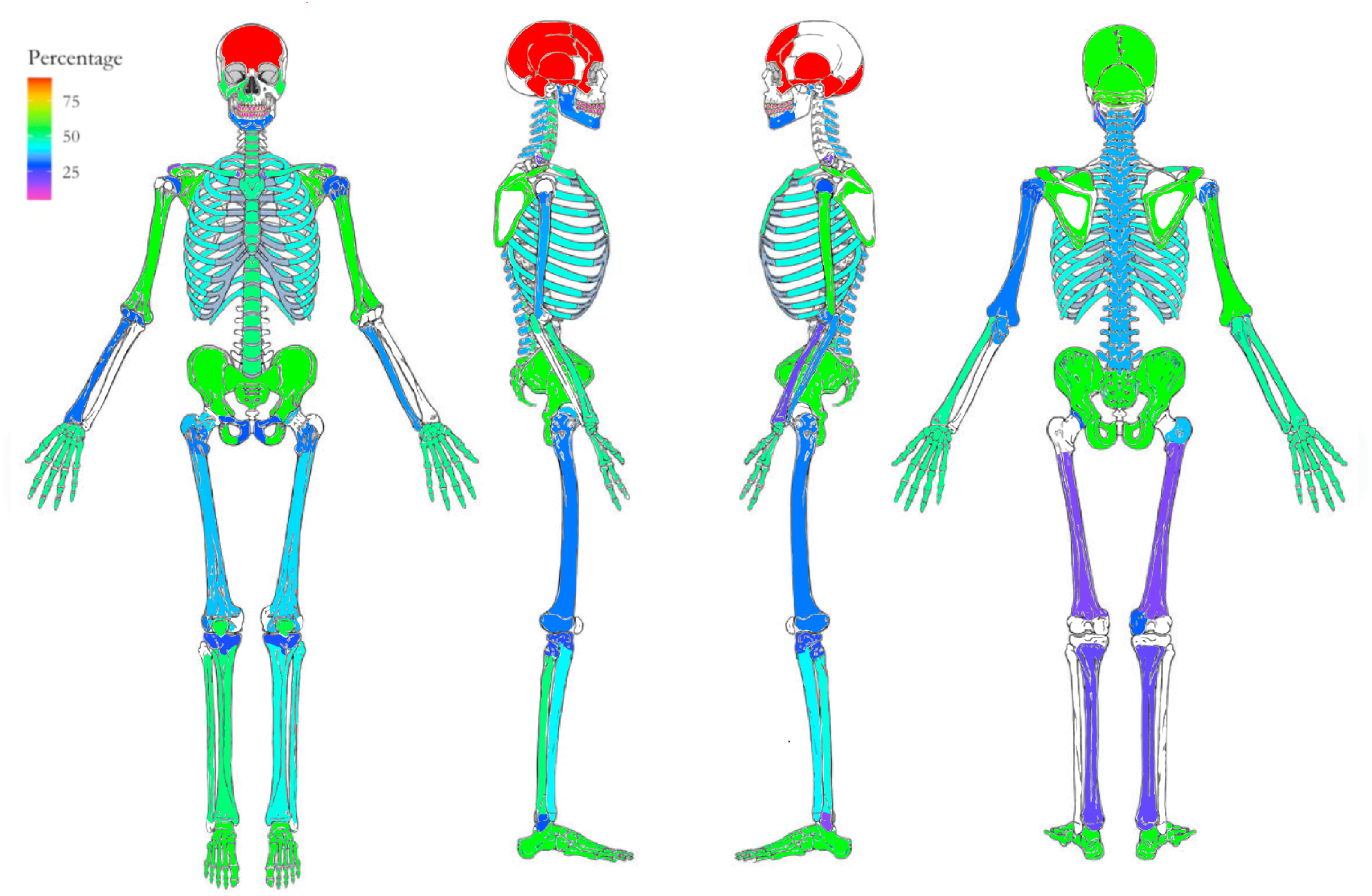
Adults percentage of individuals with pigment present, planes grouped by factor. Significantly higher quantities of ochre on the calotte of the skull. The posterior and anterior upper limbs and pelvis, have higher quantities of pigment than the lower limb.

## Discussion

This work has presented a new and accessible methodological framework by which bone surface taphonomy can be investigated free of the costs associated with bone scanning, having broader implications for osteology and forensic research. The spatial and semi-quantitative mapping of surface alteration on each individual bone enables a detailed comparison of the individuals, which can be considered against factors relating to a given research question. Breaking the surface of the adult skeleton down into 521 bone faces enables a comprehensive view of bone surface changes. When put into practice the preservation of the remains from Khok Phanom Di meant that underrepresented faces were removed, reducing the total to 424. While it could be argued that the removal of these bone faces could be a methodological adjustment, they are population specific. The use of 521 bone faces captures the maximum amount of information, which forms a starting point from which a dataset can be accessed. By splitting bones into their separate faces, distinctions can be made between a bone that has heavy ochre staining on one part but minimal or no pigment elsewhere (Figure 9). On establishing the level of preservation of an assemblage and representation of each bone face each dataset can be adjusted accordingly, as was the case with Khok Phanom Di. The use of skeletal region, bone type, anatomical plane, and side of the body, associated with each bone face enable the categories by which the bone faces can be grouped, and statistical analysis undertaken. The evidence from Khok Phanom Di suggests that simple presence or absence per bone face captures a significant level of detail of bone staining. Unidentifiable skeletal elements were excluded and bones such as juvenile epiphyses were routinely excluded. If this method is to be repeated on other assemblages, comparisons between sites must acknowledge there is a bias based on the skill of the osteologist. This limitation is not detrimental but creates similar issues to preservation/collection biases. Other considerations are the impact of comingled assemblages. The research questions that this method was devised to answer considered the relationship between surface staining and osteobiographies. If individuals cannot be identified any relationship between secondary data, such as sex or overall pathology, would be more limited. Individually-based osteobiographies could not be compared; however, specific diagnostic bones (either for sex or pathology) could be considered. Anatomical comparative assessment would be the most informative on comingled assemblages.

**Figure 9.**
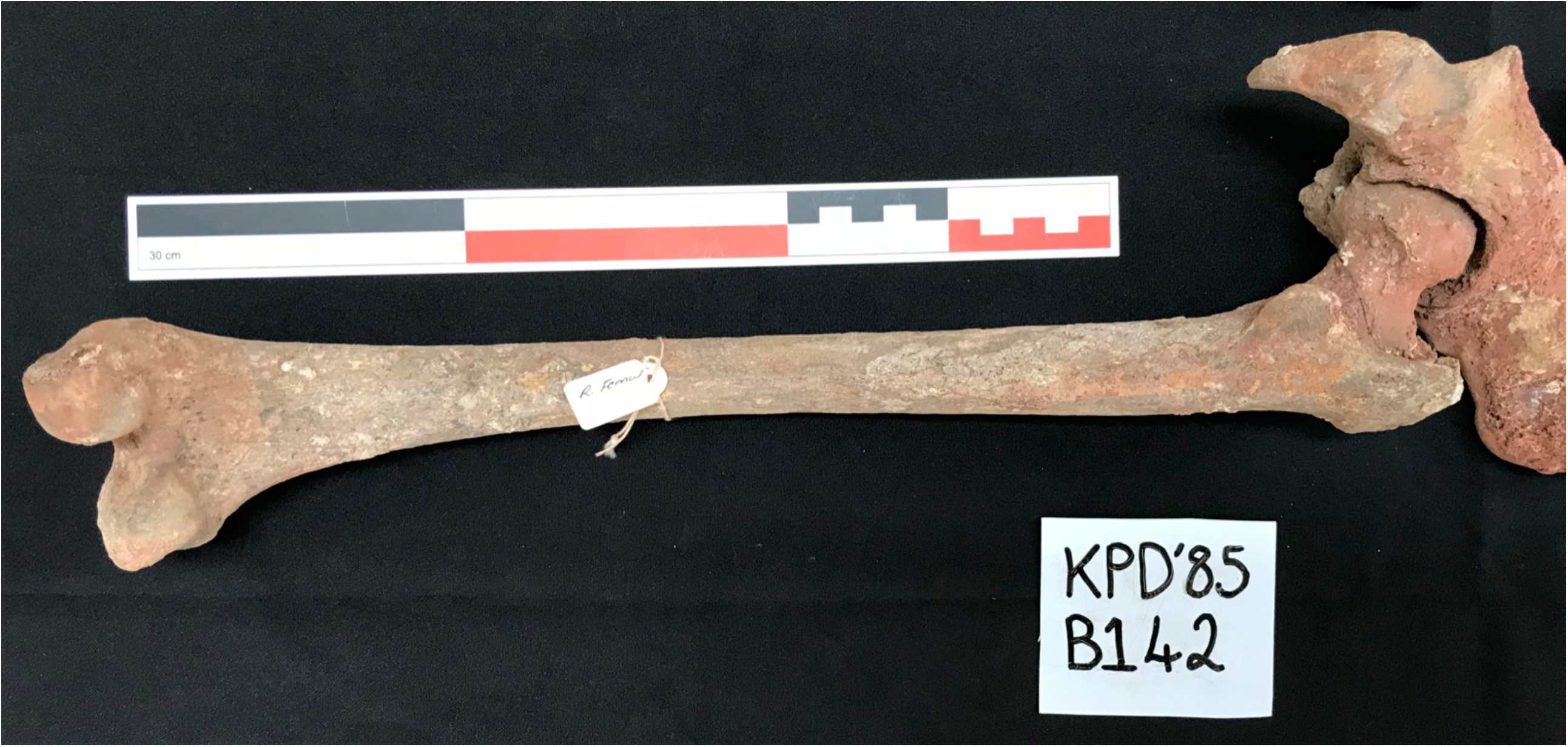
The right femur (and partial right pelvis) of B142. The femur has considerably more ochre at the proximal end, with the posterior neck, whereas the posterior shaft has comparatively little pigment. By breaking the skeleton down into bone faces, it enables these distinctions to be made and patterns compared across, and between assemblages.

## Conclusion

The results from Khok Phanom Di have demonstrated the wealth of information that can be gained from employing this methodological approach. The ease of recording and lack of equipment required (and consequently the costs) make it an extremely accessible method to osteologists, the only prerequisite being knowledge of skeletal anatomy. This advances the standardisation of bone surface recording on human remains as part of osteological protocol where taphonomic processes have occurred. The scale of recording will ultimately come down to the aims of skeletal analysis and time afforded to taphonomy as part of that. The hope is that this research has demonstrated the value and potential that bone surface mapping has to contribute to better understanding of a given site and culture, providing a practical alternative to photogrammetry.

## Supporting information

Paris and Foley recording sheet

## Acknowledgements

We express sincere gratitude to The National Fine Arts Department and the Regional Office in Nakhon Ratchasima for permission to access the human remains from Khok Phanom Di, especially Janaung Supassy and Jaruk Wilaikeaw. Further thanks to Professor Robert Foley for his supervision; and Professor Charles Higham and Dr Rachanie Thosarat who facilitated my work in Thailand. This research was carried out under the research permit of the University of Otago and supported by University of Cambridge fieldwork funding from the McDonald Institute, Department of Archaeology, and St Catharine’s College.

